# Automation workflow for high-throughput arrayed plasmid DNA preparation and quantification

**DOI:** 10.64898/2025.12.13.694144

**Authors:** Chih-Cheng Yang, Aniruddha J. Deshpande, Michael Jackson, Peter D. Adams, Jiang-An Yin, Yancheng Wu, Christopher J. Knuff, Abdullah Ghias, Anna Beketova, Chun-Teng Huang

## Abstract

High-throughput generation of arrayed plasmid DNA library using commercially available miniprep kits remains labor-intensive and costly. The yield and quality of plasmid preparations directly affect downstream applications, including arrayed viral library production and CRISPR library screening. Insufficient plasmid yield or DNA concentration often requires repeated preparations or additional DNA concentration steps to obtain adequate quantities. Similarly, higher variations in yield or quality across wells or plates can render an entire library unsuitable for subsequent experiments. To increase productivity and mitigate human intervention and errors, the present study established an automated workflow for high-throughput plasmid DNA preparation and quantification. The workflow was carried out by the Biomek i7 Hybrid automated workstation, synergizing a robotic liquid handler and multiple peripheral instruments to produce and measure plasmid DNA in a 96-well plate format. Bacterial competent cells were alkaline lysed and plasmid DNA was purified using magnetic beads, followed by quantification with the PicoGreen assay. The PicoGreen assay reported median and average yields of approximately 9.5 and 10 µg per sample, respectively, which are equivalent to 7.6 and 8 µg/mL of bacterial culture. Plasmid DNA concentrations measured by the PicoFluor fluorometer were consistently lower than those obtained using the NanoDrop UV spectrophotometer. The comparison demonstrated robust positive correlation between PicoGreen assay and NanoDrop measurements (R^2^ > 0.8). Among 480 plasmid DNA samples, average and median yields measured by the NanoDrop reached approximately 24 and 25 µg per sample per well, corresponding to 19 and 20 µg/mL of bacterial culture. Over 98% of samples exceeded the high-yield threshold of 10 µg/mL of culture, with high plasmid quality validated through DNA gel electrophoresis. Collectively, this study demonstrated a robust, scalable, and cost-effective automation platform for high throughput arrayed plasmid library generation and quantification.

## Introduction

Arrayed CRISPR library screening is a powerful functional genomics approach to identify potential therapeutic targets and their underlying mechanisms in disease models. High throughput plasmid DNA preparation (miniprep) and quantification are essential prerequisites for arrayed lentivirus production and subsequent CRISPR library screening. To replenish arrayed lentiviral libraries, lentivector plasmids must be amplified from bacterial glycerol stocks and extracted from bacterial cultures. The materials and methods for bacterial pellet resuspension, lysis, and neutralization are generally consistent across various commercially available kits [1, 2]. From the neutralized DNA lysates, several high-throughput plasmid purification methods are available, including silica membrane-based spin columns (QIAGEN), chaotropic resin-based PhyTip pipette tips (Biostage), and others. To support arrayed CRISPR DNA library preparation, we adopted a previously plasmid preparation procedure for generating the T. gonfio quadruple-gRNA DNA library, as described by Yin *et al.* at the University of Zurich, Switzerland [3], and optimized conditions suitable for automated protocols using the Biomek i7 Hybrid automated workstation to streamline the high-throughput workflow for arrayed plasmid miniprep and quantification. Reagents, including nutrient broth for bacterial culture; resuspension, lysis, and neutralization buffers for plasmid DNA isolation; and magnetic buffer for plasmid purification, were all prepared in-house. For DNA purification, we adopted a magnetic bead-based method using coated microbeads (Cytiva) to improve recovery efficiency and reduce cost. We also established an ultra-sensitive and highly reproducible fluorescence-based PicoGreen assay (Promega) for high-throughput quantification of double-stranded DNA using the Biomek i7 and Pherastar FS plate reader (BMG Labtech).

The Biomek liquid handling platform from Beckman Coulter has been applied to a wide range of automated workflows. The nucleic acid-related applications include amplicon library preparation [4, 5], molecular cloning [6], bacterial culture [7, 8], plasmid DNA purification [9], nucleic acid extraction [10], PCR and qPCR [6, 10, 11, 12, 13], mRNA and ChIP-Seq library preparation [14, 15, 16], as well as genomic workflows [17, 18, 19]. Previously, we utilized Biomek i7 Hybrid automated workstation to establish high throughput arrayed mammalian cell line cultivation [20], lentivirus production and titration [21], as well as CRISPRa library screening [22]. The present study further leveraged our Biomek i7 Hybrid liquid handler, integrated with multiple instruments, to automate a high throughput workflow for bacterial culture inoculation, plasmid DNA isolation, purification, and quantification. The plasmid yield and purity produced from these automated pipelines are reproducible and comparable to the conventional manual miniprep.

The automation workflow for high throughput arrayed plasmid DNA preparation and quantification on the Biomek i7 platform was established using the Beckman Coulter software suite. While basic liquid handling steps can be programmed using either the Biomek 5 software or SAMI EX software, we primarily utilized SAMI EX (v5.0) due to its ability to enable communication between the Biomek liquid handler and integrated peripheral instruments, as well as its flexible command interface for planning, executing, and tracking the entire workflow. The dynamic scheduler within the SAMI EX interface enables efficient time and resource management by coordinating hardware movement sequences and allowing multiple methods to run in parallel. Similar to the Biomek 5 software, SAMI EX also offers simulation capabilities to validate functionality, optimize procedures, and detect potential errors prior to execution. This robust configuration enabled fully automated, high-throughput workflows optimized for arrayed plasmid miniprep, encompassing the entire process from bacterial culture inoculation to plasmid purification and DNA quantification in a 96-well plate format.

## Materials and Methods

### 1. System configuration of the Biomek i7 Hybrid automated workstation with integrated instruments

The Biomek i7 automated liquid handling system (Product No. B87585, Beckman Coulter) is equipped with hybrid multichannel pods, a dual-pod system combining 96/384-channel heads and eight independent pipette heads, along with a barcode reader, two track grippers, a plate shuttle station, a Biotek 405 LSHTV Washer (Agilent), a BioShake 3000-T ELM microplate shaker (QInstruments), a Cytomat 2C hotel incubator (Thermo Fisher Scientific), a CloneSelect imager (Molecular Devices), and a microcentrifuge (Agilent). These components work synergistically to streamline high-throughput automation within a sterile enclosure equipped with HEPA-filtered fans. The Beckman Coulter software suite includes Biomek software 5 for method and workflow development, as well as SAMI software for centralized workflow control, scheduling, and extended method design.

### 2. Plasmid DNA preparation and quantification

The high-throughput arrayed plasmid DNA preparation (miniprep) and quantification workflow had four automated pipelines (Figure 1A): 1) bacterial culture inoculation, 2) plasmid DNA isolation, 3) plasmid DNA purification, and 4) plasmid DNA quantification. Plasmid DNA samples (pYJA5 plasmid backbone, approximately 9.1 kb) used in this study were derived from the *T. gonfio* quadruple-gRNA DNA library [3].

**Figure 1:**
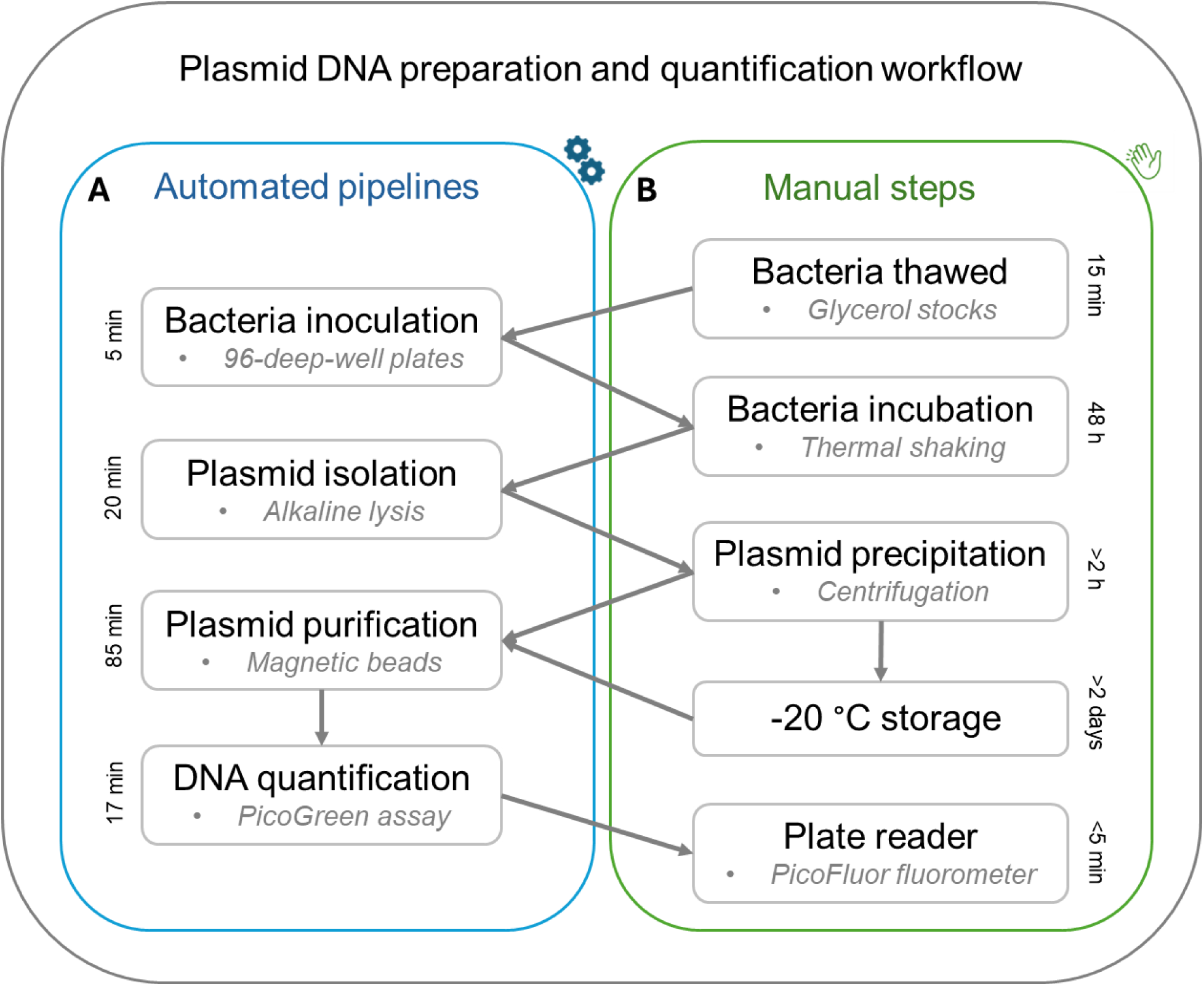
High-throughput automated arrayed plasmid DNA preparation and quantification workflow. Schematic overview of the workflow comprising four automated pipelines and four manual procedures described in Materials and Methods. **A.** Automated pipelines employed a 96-channel pipette head for liquid handling to perform multiple steps, including inoculating bacteria (Method 2.1 and Figure 2) from a glycerol stock plate into a new 96-deep-well plate, dispensing resuspension, lysis, and neutralization buffers for plasmid isolation (Method 4 and Figure 3), adding 100% ethanol for DNA precipitation, purifying DNA using magnetic beads with 70% ethanol washes and TE buffer resuspension (Method 6 and Figure 4), and diluting plasmid DNA with PicoGreen reagents for fluorescence-based DNA quantification (Method 7 and Figure 5). **B.** Procedures that required manual processing included thawing bacterial plates (Method 2.1), removing and sealing plates, transferring inoculated plates to thermal shakers for 48-hour culture (Method 3), moving plates in and out of the freezer for short- or long-term storage (Method 5), and loading plates into the plate reader for fluorescence analysis (Method 7).

#### 2.1. Bacterial culture inoculation

The automation deck layout was shown in Figure 2E. The maximum capacity for this workflow is four 96-well plates per set (Figure 2B, Table 1). A set of four 96-well plates containing freshly thawed arrayed bacterial glycerol stocks was centrifuged at 500 g-force for 1 min before removing the seal. A volume of 25 µL of bacteria from each well was transferred into four new 96-deep well plates with 1.2mL of fresh Terrific Broth (12 g/L Bacto tryptone, 24 g/L Bacto yeast extract, 0.4% Glycerol, 0.017 M KH_2_PO_4_, and 0.072 M K_2_HPO_4_) with 15 mg/L trimethoprim and 15 mg/L tetracycline antibiotics, followed by pipetting up-and-down at 100 μL/s speed for 6 cycles using a 96-channel pipette head.

**Figure 2:**
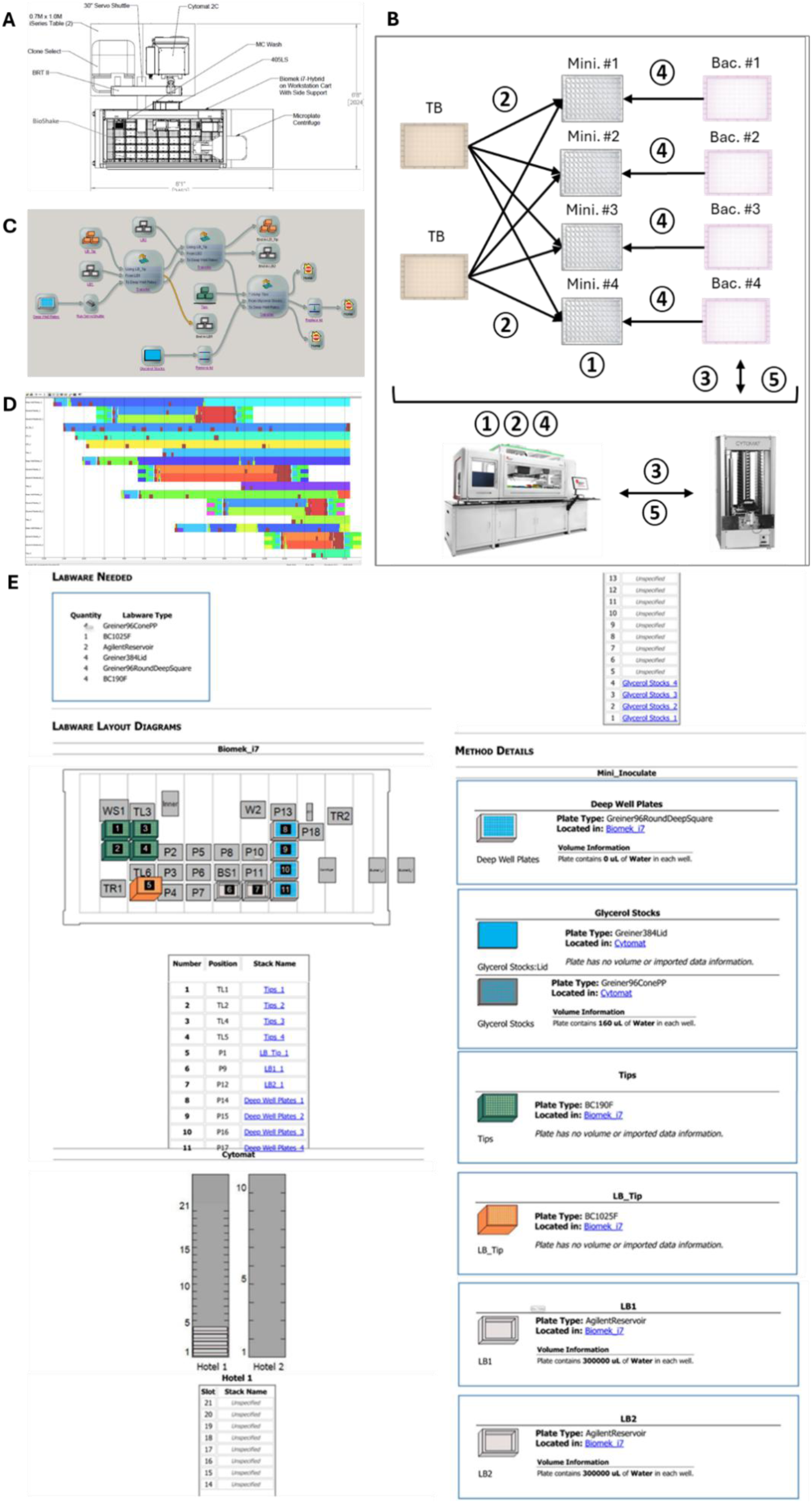
High-throughput pipeline for automated arrayed bacterial inoculation. **A.** Hardware layout of the Biomek i7 Hybrid automated workstation. **B.** Schematic overview of arrayed bacterial inoculation. ① Four empty 96-deep-well plates were initially placed on the Biomek deck. ② 1.2 mL of terrific broth (TB) were transferred from reservoirs to the 96-deep-well plates using the 96-channel pipette head. ③ Four 96-well round-bottom plates containing freshly thawed bacteria glycerol stocks were transferred to the Biomek i7 deck via the servo shuttle and track gripper. The lids were removed and placed on the deck by the gripper. ④ 50 µL of bacteria glycerol solution was transferred from the round-bottom plates to the deep-well plates and mixed thoroughly using the 96-channel pipette head. ⑤ The lids were replaced, and the plates containing leftover bacterial glycerol stocks were returned to the incubator. **C.** Pipeline overview under the SAMI EX interface. SAMI EX software was utilized for the method creation and pipeline design. **D.** SAMI’s scheduling and time estimation for the pipeline illustrated the chronological timestamps of each labware. **E.** The labware setup report provided a reference point and summarizes the SAMI EX deck layout, labware, and associated conditions.

**Table 1.** Time required for each automation pipeline.

|  | 1 plate | 2 plates | 3 plates | 4 plates |
| --- | --- | --- | --- | --- |
| **Bacterial culture inoculation pipeline.** | 5 min | 7.5 min | 11 min | 16 min |
| **Plasmid DNA isolation pipeline.** | 20 min | N/A | N/A | N/A |
| **Plasmid DNA purification pipeline.** | 85 min | N/A | N/A | N/A |
| **Plasmid DNA quantification pipeline.** | 17 min | N/A | N/A | N/A |

N/A indicates 'not available' for the desired plate quantity due to protocol design, limited deck capacity or the restricted incubation window of sample and reagent.

### 3. Bacteria culture incubation

After bacterial inoculation, four bacterial deep-well plates were manually removed out of the deck, sealed with breathable AeraSeal film (Cat# A9224-50EA, Excel Scientific), and incubated on a TS-300 incubator shaker (Pioway Medical) at 900 rpm at 30°C (Figure 1B). After 48 h of incubation, the bacterial culture plate was spun down one at a time via Allegra X-15R (Beckman Coulter) at 4,000 g for 15 min. Processing of the second, third, and fourth plates was postponed until the preceding plate had completed the plasmid DNA precipitation step to avoid scheduling conflicts and accommodate the limited liquid-handling capacity. After centrifugation and manual removal of the seal, the plate was manually inverted to discard the supernatant and placed upside-down on paper towels to remove any remaining liquid. The plate containing dried bacterial pellets was manually placed on the designated deck position.

### 4. Plasmid DNA isolation

The automation layout of the deck was shown in Figures 3D and 3F. This pipeline is designed for processing one plate at a time. Two hundred microliters of resuspension (RES) buffer (50 mM Tris-HCl, PH 8.0, 10 mM EDTA, 100 mM glucose, and 100 mg/L RNaseA) was dispensed into each well of a 96-well plate containing bacterial pellets. One plate was then placed on a BioShake 3000-T ELM microplate shaker to vortex at 1,500 rpm for 5 min, followed by pipetting up and down using a 96-channel pipette head until the bacterial pellets were completely resuspended. The bacterial suspension was lysed with 200 µL of lysis (LYS) buffer (1% SDS and 0.2 N NaOH), mixed by pipetting a volume of 400 µL six times at 500 μL/s speed (tip positioned 0.2 mm above the well bottom), and incubated for 5 min. The entire alkaline lysis step should not exceed 5 min. The lysed bacterial suspension was then neutralized with 200 µL of neutralization (NEU) buffer (3 M potassium acetate and 11.5% glacial acetic acid, PH 5.5), and mixed by pipetting a volume of 600 µL twelve times at 400 μL /s speed, with the tip positioned 0.2 mm above the well bottom.

**Figure 3:**
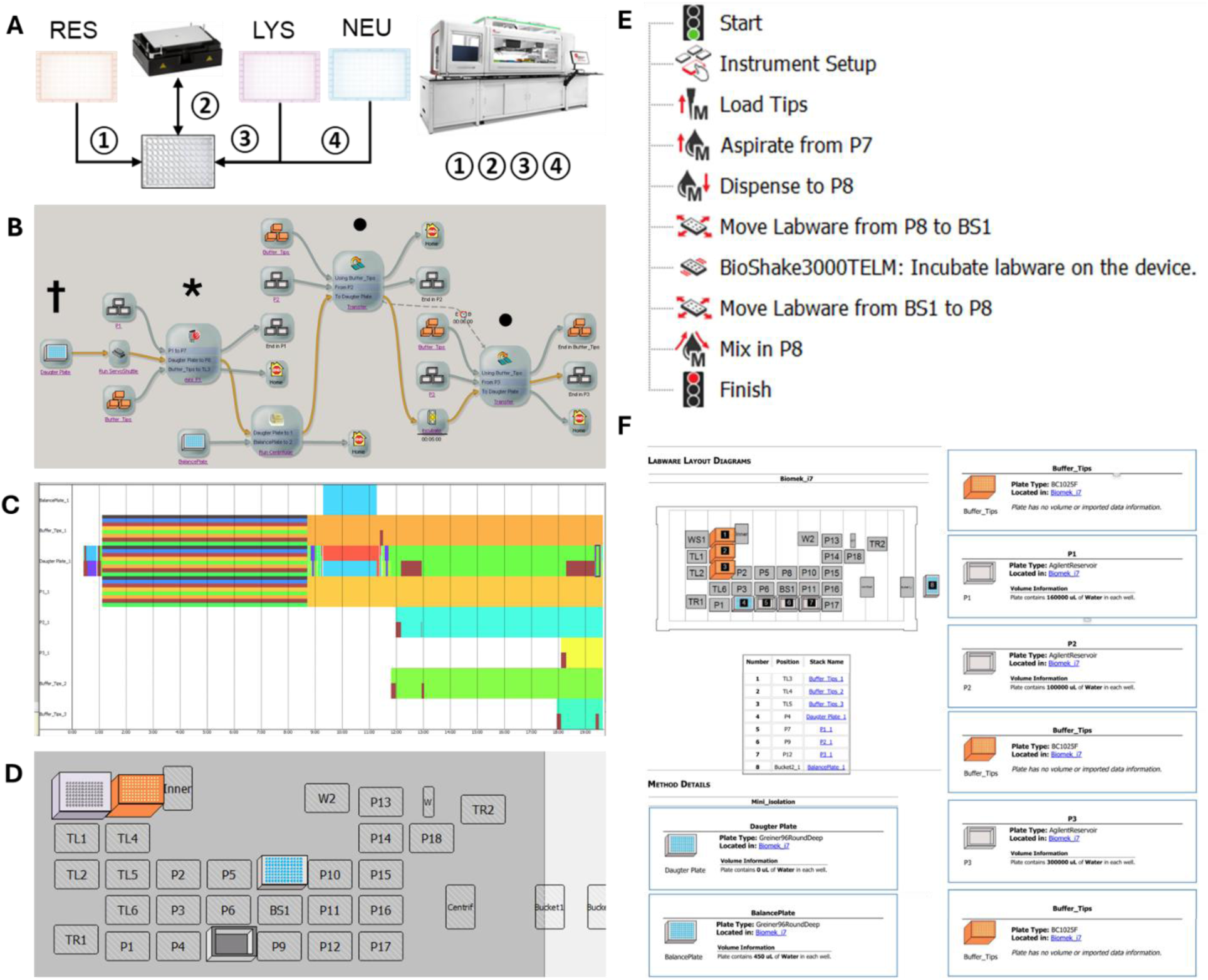
High-throughput pipeline for automated arrayed plasmid DNA isolation. **A.** Schematic overview of arrayed plasmid isolation using Biomek i7 liquid handler and BioShake. ① RES buffer was transferred from a reservoir to a 96-deep-well plate containing dry bacterial pellets on the Biomek deck. ② The plate was moved to the shaker by the gripper for vortexing, followed by pipetting up and down using the 96-channel pipette head to fully resuspend the bacteria. ③ LYS buffer was then transferred from a reservoir to the plate, followed by pipetting up and down to ensure complete bacterial lysis. ④ NEU buffer was added from a reservoir to the plate, followed by pipetting up and down to neutralize the alkaline lysate. Biomek 5 method node: asterisk symbol; SAMI EX method node: black dot symbol. **B.** Pipeline overview under the SAMI EX interface. Both SAMI EX and Biomek 5 software were utilized for the method development. **C.** Scheduling and time estimation of the entire pipeline in SAMI EX, showing the chronological timestamps of each labware item. **D.** Deck layout for RES buffer transfer and plate vortexing, indicated by the asterisk in Figure 3B. **E.** Biomek 5 method for RES buffer transfer and plate vortexing (asterisk in Figure 3B). **F.** Labware setup report, indicated by the dagger symbol in Figure 3B, providing an overview of the SAMI EX deck layout, including labware details and conditions.

### 5. Plasmid DNA precipitation

After mixing neutralized suspension, the plate was manually removed from the deck, sealed with polyolefin transparent sealing tape/film (Ref# 95.1994, Sarstedt Inc), and centrifuged at 4,000 g for 15 min to precipitate debris. After centrifugation, the plate was relocated back to the incubator and 400 µL of supernatant was transferred from the neutralized plate into new 96-deep well plates at 50 µL/s speed, with the tip positioned 10 mm above the well bottom (automation protocol not shown). It is optional to pause the automation and ensure uniform volume across all wells before proceeding to the ethanol precipitation step. To precipitate the plasmid DNA, 1 mL of 100% molecular-grade ethanol was added to each well and mixed by pipetting up and down six times at 100 µL/s speed (automation protocol not shown). The DNA-precipitated plate was manually removed out of the deck, sealed, and incubated at −20°C overnight. Alternatively, the plate could be processed immediately if the ethanol was pre-chilled to −20°C. The plate may also be stored at −20 °C for several hours to days before proceeding with downstream steps. Next, the DNA-precipitated plate was centrifuged at 4,000 g for 15 min at 4°C. After centrifugation, the seal was removed by hand and the supernatant was discarded by manually inverting the plates, which were then placed upside-down on paper towels to absorb any residual liquid.

### 6. Plasmid DNA purification

The plate containing dried DNA pellets was transferred back to the Biomek deck and placed on the microplate shaker for vortexing at 1,500 rpm for 6 min to fully dissolve the DNA after 50 µL of molecular-grade water was dispensed into each well. After DNA resuspension, 75 µL of beads buffer (2.5M NaCl, 0.2 kg/L PEG 8000, 0.05% Tween 20, 10 mM Tris base, 0.336% 1 N HCl, and 1mM EDTA) and 50 µL of a 1:50 dilution of SpeedBeads magnetic carboxylate-modified particles (Millipore Sigma, Cat. #GE65152105050250) in beads buffer were added to each well, followed by vortexing at 1,000 rpm for 10 min using the microplate shaker to facilitate DNA-bead complex formation. The plate containing DNA–bead complex solution was then incubated on a magnet rack (homemade, base 3D-printed and integrated with neodymium disc magnets, N55, 0.125 in × 0.375 in, Part #N55P125375) for 10 min to magnetically separate the beads from the liquid. After the DNA–bead complexes had attached to the walls of the wells, the liquid was slowly removed using a 96-channel pipette head at 100 μL/s speed, with the tip positioned 1 mm above the well bottom, while the plate remained on the magnet rack. One milliliter of 70% ethanol was added (at 100 μL/s speed, with the tip positioned 6 mm below the liquid surface), incubated for 5 min, and then removed (at 100 μL/s speed, with the tip positioned −11 mm from the well top and select “follow liquid level” when aspirating). The entire wash procedure was repeated twice per well to clean the DNA-bead complexes on the magnet rack. After the second wash, an additional aspiration was performed to remove any remaining liquid, and the DNA–bead complexes were then incubated for 15 min to air-dry. After being transferred from the magnet rack to the microplate shaker, the plate received 150 μL of pre-warmed TE buffer (65 °C) to resuspend the DNA from the bead aggregates by vortexing at 65 °C for 15 min. After a homogeneous suspension was formed, the plate was transferred back to the magnet rack and incubated for 10 min to magnetically separate the beads from the DNA solution. Approximately 150 μL of DNA solution per well was transferred to a new 96-well conical-bottom plate on a magnet rack (homemade, flat base with a layer of neodymium bar magnets, N52, 60 x 10 x 3 mm, ASIN # B0D5MFMT9K, DIYMAG) and incubated for 5 min for a second bead clarification step. The final clear solution containing purified plasmid DNA was then transferred to a new 96-well conical-bottom plate for downstream quantification and experiments.

### 7. Plasmid DNA quantification

After plasmid purification, the DNA concentration of each sample in the 96-well conical-bottom plate was measured using the PicoGreen assay (Life Tech). To generate a standard curve within the linear detection range, lambda DNA was used as the standard control and prepared starting at 2 ng/mL through six 1:3 serial dilutions in the first column of the 96-well reagent plate, with 120 µL per well (final volumes), using 8 independent pipette heads. The last well (the H1 position) of the first column, containing 100 µL of TE buffer only, served as the negative control. Columns 2-4 of the reagent plate contained 200 µL of PicoGreen reagent (1:200 dilution with TE). The quadruple-1 (Q1) well positions in columns 1, 3, and 5 and rows A, C, E, G, I, K, M, and O of a new 384-well assay plate were first filled with 20 µL of serially diluted positive-control lambda DNA standards and negative-control TE buffer from the first column of the 96-well reagent plate (with triplicate wells per control) using 8 independent pipette heads.

Four microliters of each plasmid DNA sample from a 96-well conical-bottom plate were transferred into another 96-well conical-bottom plate containing 76 µL of TE buffer, supplied from the reservoir, to generate an intermediate 1:20 dilution stock. Subsequently, 10 µL of the intermediately diluted DNA samples were transferred into a new 96-well conical-bottom plate pre-filled with 190 µL of TE buffer from the reservoir to achieve a total 1:400 dilution relative to the original DNA sample. Lastly, 20 µL of DNA solution from the 1:400 dilution 96-well plate were transferred to the quadruple-3 (Q3) and quadruple-4 (Q4) well positions of the 384-well assay plate, which served as duplicate wells for the assay readout of each sample. All transfers were performed using the 96-channel pipette head.

Twenty microliters of PicoGreen reagent were transferred from the reagent plate into the Q1, Q3, and Q4 well positions of the assay plate using 8 independent pipette heads. The assay plate underwent a brief centrifugation and BioShake vortexing. After a 10-minute incubation at room temperature, fluorescence signals were measured using the PHERAstar FS plate reader (BMG Labtech).

## Result

### High-throughput automation design

The Biomek i7 Hybrid automated workstation is a dual-pod liquid handling system which enabled high throughput automated arrayed plasmid preparation workflow in this study. The system had a left pod equipped with interchangeable 96 or 384-channel pipette heads for microplate stamping, and a right pod incorporated with eight independent pipette heads for cherry-picking and plate reformatting. In addition to the liquid handler, multiple integrated instruments contributed specific functions required for the workflow (Figure 2A). The Cytomat 2C incubated cell culture in multiple microplate formats. The CSI enabled cell confluency monitoring at high throughput. The BioShake 3000-T ELM shaker supported heating and vortexing capabilities, while the microcentrifuge provided rapid clarification of supernatants.

### Bacterial culture inoculation pipeline

Within the plasmid miniprep workflow using a 96-deep-well plate format, the first pipeline for bacteria inoculation was optimized from one to four plates, with each sample per well containing 1.2 mL of plasmid-transformed bacterial culture, illustrated in Figure 2B. SAMI EX software was used to develop methods and designed the pipeline. Its interface simulated, scheduled, and executed pipelines (Figure 2C) by managing hardware multitasking and labware movements in a chronological, timestamp-based manner (Figure 2D). The labware setup report provided an overview of the SAMI deck layout, including details of the labware and reagents (Figure 2E). The runtime for this bacteria inoculation pipeline to prepare one to four 96-well plates was 5 to 16 minutes (Figure 2D, Table 1).

### Plasmid DNA isolation pipeline

Plasmid DNA was first isolated from bacterial pellets using the alkaline lysis method. To mitigate liquid-handling capacity limitations, which could lead to scheduling conflicts and extended waiting times, the plasmid isolation pipeline was optimized to process one 96-well plate per run (Figure 3A). Thus, sequential runs of this pipeline were employed to accommodate multiple plates generated from the preceding bacterial inoculation pipeline. Methods developed using Biomek 5 and SAMI EX software programs were integrated into a single automated pipeline that was controlled, scheduled, and executed through the SAMI EX interface. (Figure 3B). SAMI EX estimated the runtime for processing one plate in this pipeline was 20 minutes (Figure 3C and Table 1). Biomek 5 software was utilized to coordinate complex steps, including buffer transfer, plate movement, BioShake operation, and suspension mixing, under a distinct deck layout (asterisk symbol in Figure 3B; Figures 3D and 3E). In contrast, SAMI EX was used to develop general procedures such as multiple buffer transfers and its labware setup report summarized the initial deck layout and labware details (dagger symbol in Figure 2B; Figure 3F).

### Plasmid DNA purification pipeline

The pipeline starts with resuspending semi-dry DNA pellets in molecular-grade water in a 96-deep-well plate. These pellets were produced through the plasmid DNA isolation pipeline and subsequently subjected to ethanol precipitation, centrifugation, and supernatant removal manually. Plasmid DNA was then purified via automated liquid handling using magnetic bead affinity binding, ethanol washes, and TE buffer elution performed on a custom 3D-printed magnetic plate (Figure 4A). Both SAMI EX and Biomek 5 contributed to the pipeline development (Figure 4B), which required 85 minutes to process one 96-well plate per run according to SAMI EX scheduling and time estimation (Figure 4C and Table 1). This pipeline consisted of three Biomek 5 nodes to perform sophisticated, repetitive mixing and liquid-transfer steps, as well as integrate the usage of BioShake and magnetics plate. (asterisk symbol in Figure 4B; Figures 4D). Starting with the labware setup report, which outlines the initial deck layout and labware details (Figure 4E), the SAMI EX node (dagger symbol in Figure 4B) coordinated instrument and executed basic pipetting tasks, including DNA pellet resuspension in water and subsequent microplate centrifugation.

**Figure 4:**
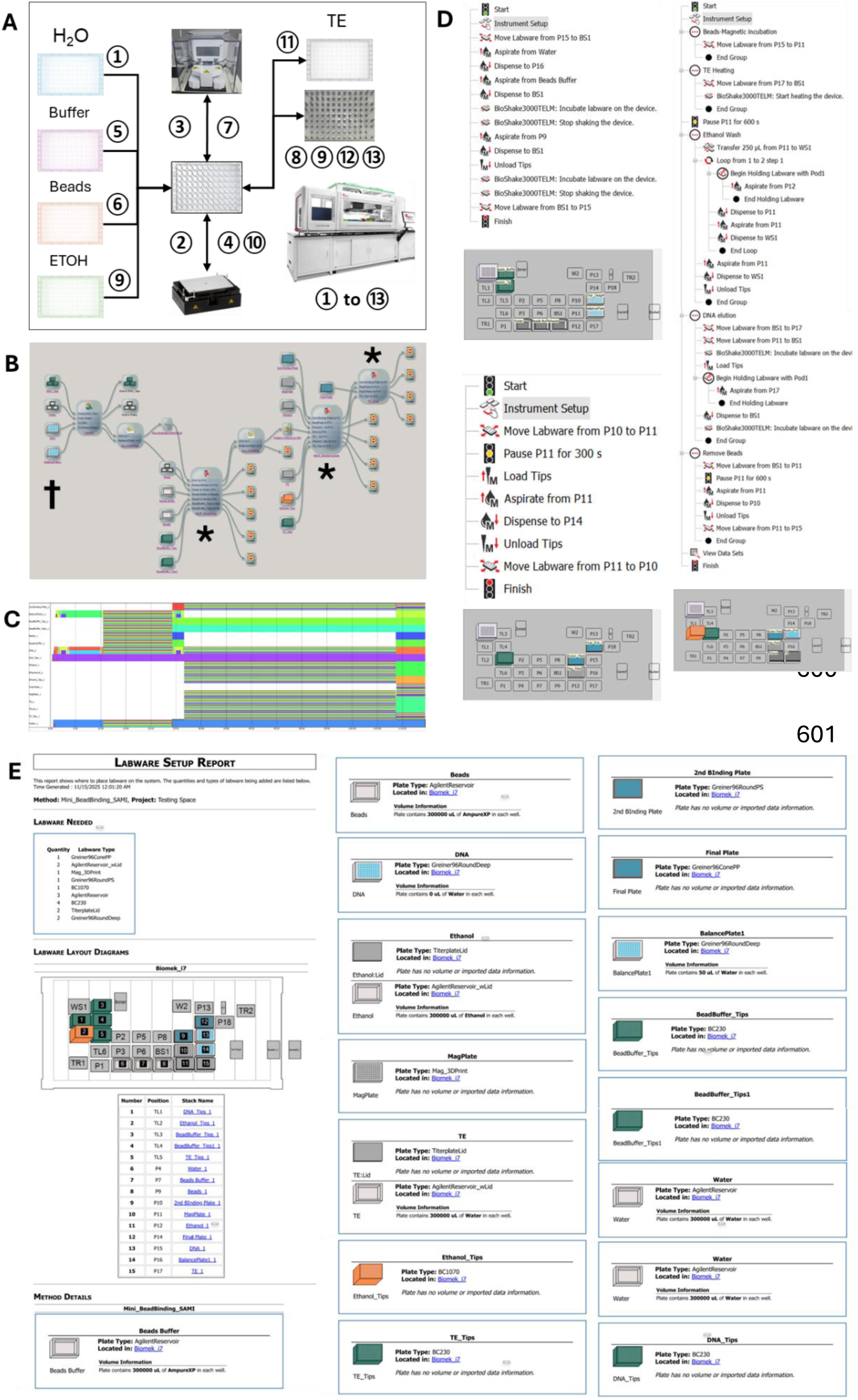
High-throughput pipeline for automated arrayed plasmid DNA purification. **A.** Schematic overview of arrayed plasmid DNA purification using the Biomek i7 liquid handler, BioShake, microplate centrifuge, and magnetic plate. All steps (① - ⑬) were performed within the Biomek i7 system. ① Molecular-grade water was added to the 96-deep-well plate containing dry DNA pellets. ② The plate was transferred to the BioShake for vortexing.③ The plate was transferred to the centrifuge to spin down DNA suspension. ④ The plate was returned to the BioShake and remained there for the next two steps. ⑤ Bead buffer was added to the plate, and then vortexed. ⑥ Bead solution was added to the plate, followed by vortexing and incubation to enable DNA-bead binding. ⑦ The plate was transferred to the centrifuge for a brief spin-down. ⑧ The plate was placed on the magnetic plate for DNA-bead isolation and remained there for the subsequent steps, except step 10. ⑨ After liquid removal by pipetting, 70% ethanol (ETOH) was added to the plate for DNA washing, followed by liquid removal and a repeat wash. ⑩ After removing the second wash, the plate was transferred from the magnetic plate to the pre-warmed BioShake. ⑪ TE buffer was added to the plate, followed by vortexing and incubation on the BioShake for DNA elution. ⑫ The plate was then placed back onto the magnetic plate for DNA-bead separation. ⑬ The supernatant containing plasmid DNA was transferred to a new 96-well conical-bottom plate, followed by a secondary transfer to another plate to remove any residual beads. **B.** Pipeline overview displayed in the SAMI EX interface. Method development and pipeline design were performed using the SAMI EX and Biomek 5 software platforms. **C.** SAMI EX provides scheduling and time estimation for the entire pipeline, displaying the chronological timestamps for each labware item. **D.** Overview and deck layout of the three Biomek 5 methods: buffer and bead treatment for DNA-bead binding (top-left panel), ethanol wash and TE addition for DNA washing and elution (right panel), and DNA solution clarification through bead removal (bottom-left panel), as indicated by the asterisk in Figure 4B. **E.** The labware setup report, marked by the dagger symbol in Figure 4B, outlines the SAMI EX deck layout and labware conditions.

### Plasmid DNA quantification pipeline

Plasmid DNA prepared at a mini-scale is typically quantified using a NanoDrop UV spectrophotometer, which is often operated manually and is not suitable for high-throughput automation. In contrast, the PicoGreen assay is specific for concentration measurement of double-stranded DNA and requires dilution of DNA samples and fluorescence detection, a procedure that can be easily adapted for automation and scaled to high-throughput formats. Therefore, we developed an automated pipeline for PicoGreen assay processing on the Biomek i7 platform to generate a 384-well assay plate ready for the PicoFluor fluorometer reading. (Figure 5A). The entire pipeline of plasmid DNA quantification is illustrated in Figure 5B and is managed and scheduled via SAMI EX (Figure 5C). Due to the extensive pipetting and serial dilution requirements, four Biomek 5 nodes were developed and integrated into the pipeline (asterisk in Figure 5B; Figure 5D). SAMI EX estimates that the pipeline, from a single 96-well DNA sample plate to a 384-well assay plate, can be completed in 17 minutes (Figure 5C and Table 1). The labware setup report (Figure 5D), marked by the dagger in Figure 5B, provides an overview of the SAMI deck layout, including the labware names, types, quantities, positions on the Biomek i7 and peripheral instruments, and associated method details.

**Figure 5:**
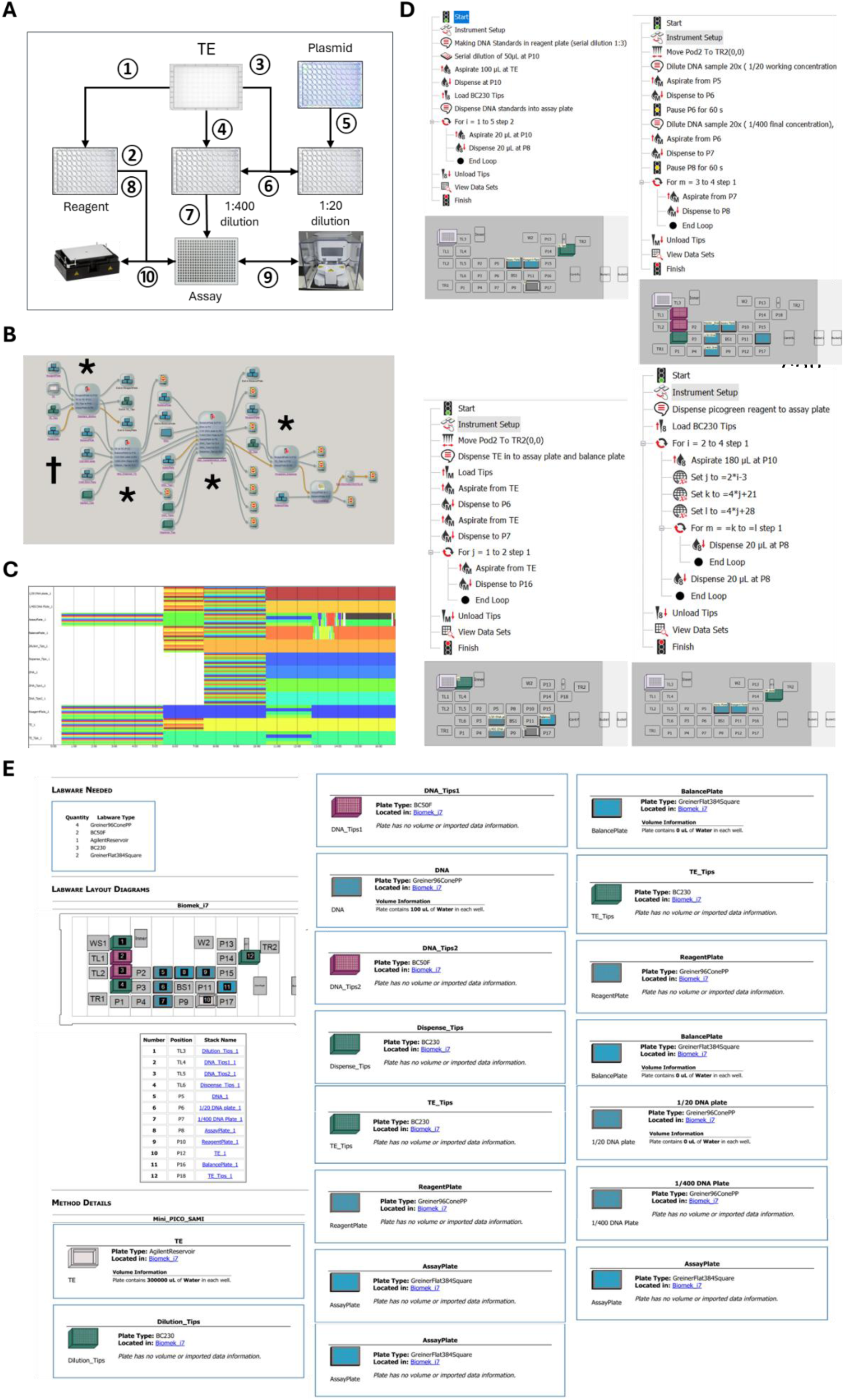
High-throughput pipeline for automated arrayed plasmid DNA quantification. **A.** Overview of arrayed plasmid DNA quantification using the Biomek i7 liquid handler and microplate centrifuge. All steps (① - ⑩) were performed within the Biomek i7 system. ① A 96-well plate containing 2 ng/µL standard DNA in well A1 and PicoGreen reagent in columns 2-4 was designated as the reagent plate. TE buffer, stored in a reservoir, was used as the diluent to perform a six-point, three-fold serial dilution from wells A1 to G1 in column 1. The negative control well, H1, contained TE buffer without DNA. ② Twenty microliters of the serially diluted DNA and TE buffer in column 1 were transferred from the reagent plate to the quadruple 1 (Q1) well positions in columns 1, 3, and 5 of a new 384-well assay plate as the standard and negative controls, each in triplicate. ③ TE buffer was transferred from the reservoir to a new 96-well conical-bottom plate (designated as the 1:20 dilution plate) to serve as the diluent for the first DNA sample dilution. ④ TE buffer was also transferred from the reservoir to another 96-well conical-bottom plate (designated as the 1:400 dilution plate) to serve as the diluent for the second dilution. ⑤ Purified plasmid DNA samples in a 96-well plate were transferred to the 1:20 dilution plate to generate an intermediate dilution stock. ⑥ The intermediately diluted DNA was transferred to the 1:400 dilution plate to achieve the final dilution. ⑦ The final diluted DNA samples from the 1:400 dilution plate were transferred to the Q3 and Q4 wells of the assay plate in duplicate. ⑧ PicoGreen reagent was transferred from the reagent plate to the Q1, Q3, and Q4 well positions of the assay plate. ⑨ The assay plate was subjected to a brief spin-down. ⑩ The assay plate was then placed on the BioShake for sample mixing prior to measurement. **B.** Pipeline overview as displayed in the SAMI EX interface. Method development and pipeline design were carried out using the SAMI EX and Biomek 5 software. **C.** SAMI EX generated a schedule and time estimate for the entire pipeline, including chronological timestamps for each labware item. **D.** Overview and deck layout of the four Biomek 5 methods: serial dilution of the standard DNA (top-left panel), spotting of the standard and negative controls (bottom-left panel), dilution of the plasmid DNA samples (top-right panel), and dispensing of the PicoGreen reagent (bottom-right panel), as indicated by the asterisk in Figure 5B. **E.** The labware setup report, marked by the dagger in Figure 5B, outlines the SAMI EX deck layout and labware conditions.

To compare different DNA quantification assays, the concentration of 480 plasmid DNA samples, equivalent to five 96-well plates, obtained from the plasmid isolation and purification pipelines were measured using both the NanoDrop and PicoGreen methods. When quantifying DNA using the NanoDrop, average and median yields reached approximately 24 and 25 µg per sample per well of 1.2 mL bacterial culture, respectively, which are equivalent to 19 and 20 µg/mL of bacteria (Figure 6A). In comparison with the NanoDrop measurements, the PicoGreen assay consistently reported lower DNA concentrations (Figure 6B), with median and average yields of approximately 9.5 and 10 µg per sample, respectively, corresponding to 7.6 and 8 µg/mL of culture (Figure 6B). On average across the interquartile range (Q1-Q3), NanoDrop readings were approximately 2- to 3-fold higher than those obtained with the PicoGreen assay (Figure 6B, data not shown). Despite this difference in absolute values, the two methods exhibited a strong positive correlation, with an R² greater than 0.8 between NanoDrop and PicoGreen assay measurements (Figure 6C). After DNA concentration normalization, high supercoiled DNA content and overall plasmid quality were confirmed by gel electrophoresis, with a representative example presented in Figure 6D.

**Figure 6:**
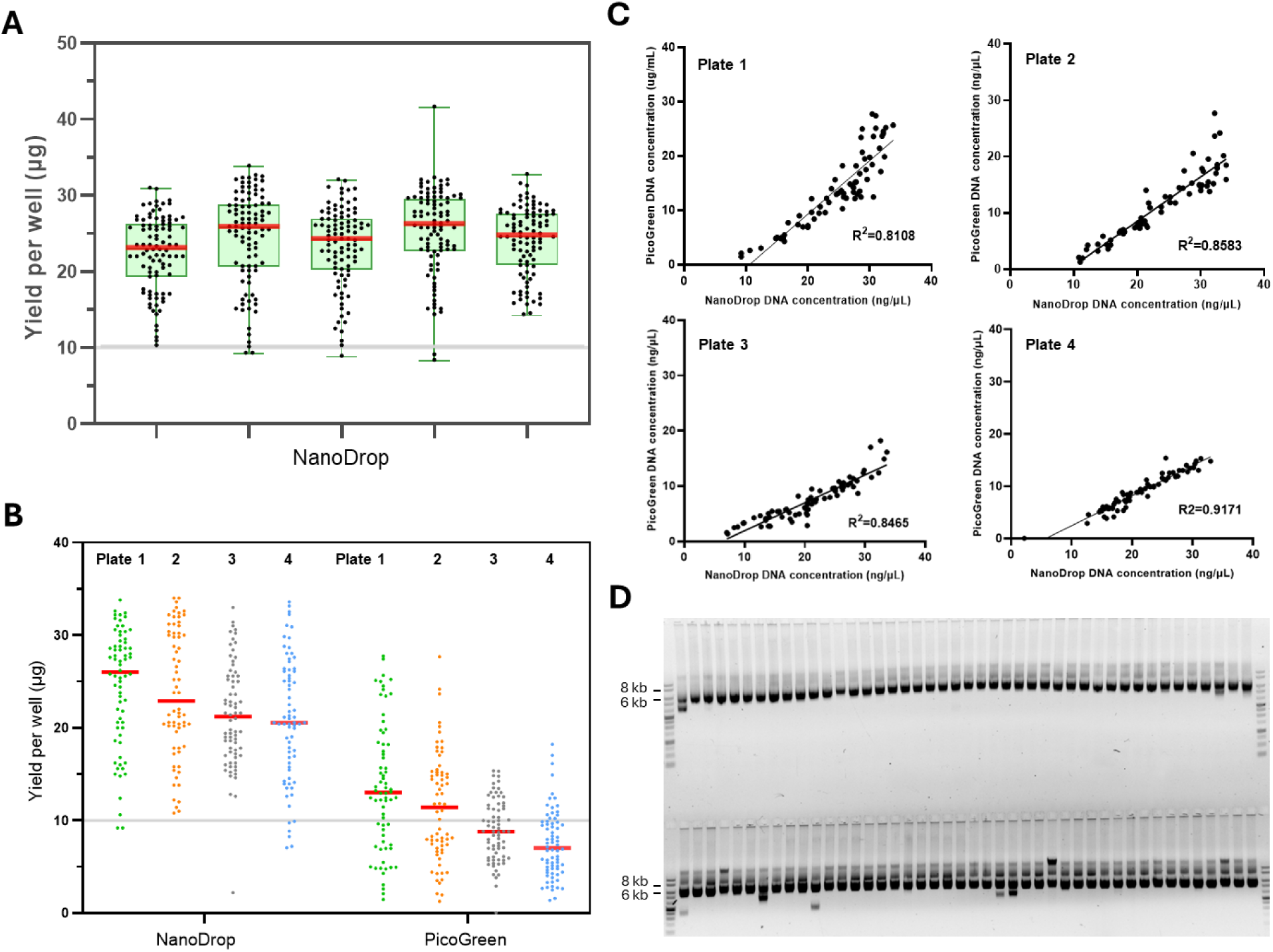
Evaluation of automated plasmid DNA miniprep and quantification workflow. **A.** Plasmid DNA quantities from five 96-well plates (480 samples from the *T. gonfio* library), generated using the Biomek i7 and measured by the NanoDrop, are presented as overlapping scatter and box plots. The median value is indicated by a red line, and the green box denotes the interquartile range. The gray solid line marks the high-titer threshold of 10 µg of DNA. **B.** Comparison of DNA concentration measurements between the NanoDrop and PicoGreen assay across four 96-well plates. The median is shown in red, while the gray solid line represents the 10 µg DNA threshold. **C.** Scatter plots depicting the correlation (R²) between DNA concentrations measured by the NanoDrop and the PicoGreen assay across four plates. The values on the X- and Y-axes represent individual DNA concentrations measured by the two methods. **D.** Gel electrophoresis image showing the migration pattern of equal amounts of plasmid DNA (60 ng per lane) after PicoGreen-based concentration normalization, alongside 1 kb DNA ladders. The intensities of the 6 kb and 8 kb ladder bands each represent 2.8 ng of DNA.

## Discussion

The workflow of plasmid minipreps in this study was validated using multiple plasmid DNA backbones, including lentiviral, retroviral, and non-viral vectors (data not shown), as well as different antibiotic selection markers such as ampicillin, trimethoprim, and tetracycline. Chemically competent *E. coli* strains Stbl3 (Cat# C737303, Thermo Fisher Scientific) and NEB Stable (Cat# C3040, New England Biolabs) were tested in this workflow. Across backbones, plates and batches, median yields were in the range of 15-30 µg per well per sample, equivalent to approximately 12-24 µg per mL of bacterial culture (data not shown), indicating the workflow is compatible with plasmids of both low and high copy numbers. Terrific Broth, rather than 2xYT or LB medium, was selected as a rich growth medium suitable for bacterial culture over 48 hours. Bacterial cultures were incubated at 30 °C with vigorous shaking at 900 rpm to minimize bacterial precipitation, cell death from overgrowth, and plasmid recombination. The alkaline lysis method was utilized to extract plasmid DNA from bacteria cultures, followed by preliminary DNA enrichment through ethanol precipitation. To minimize bacterial genomic DNA contamination, the incubation time for both lysis and neutralization steps was limited to 5 minutes. After separating the plasmid-containing supernatant from cell debris and genomic DNA/proteinaceous aggregates, an optional step to further clarify the lysate and improve purity was to manually filter the lysate through a vacuum-driven NucleoSpin 96 Plasmid Filter Plate (Cat. #740483, Takara Bio). DNA aggregation was improved by incubating at −20 °C during ethanol precipitation, which also enabled long-term storage. To further isolate plasmid DNA after initial ethanol precipitation, this study adopted a bead-based method, moving away from silica-based spin columns or similar anion exchange methods that typically require high-speed centrifugation or vacuum suction to drive flow through the column. During plasmid purification, the workflow adopted two washes of 70% ethanol as desalting steps which was much simpler compared with the washing steps required in anion-exchange-based approaches. DNA binding, washing, and elution were performed within the same deep-well microplates to minimize sample and flocculant contamination from the vacuum-based methods. The PicoGreen assay enabled sensitive detection and quantification of DNA concentrations, while minimizing fluorescence contributions from RNA and single-stranded DNA. The linear detection range, established from the standard curve using positive control DNA on the PicoFluor fluorometer, was between 1 and 1000 ng/mL. Steps such as bacteria incubation, plasmid precipitation, and PicoGreen assay analysis, required manual processing rather than full automation due to limitations of the current automation system, including the restricted speed of the thermal shaker and microplate centrifuge, lack of an integrated refrigeration unit, absence of automatic plate sealer and seal remover, and the non-integration of a plate reader.

For the software, although SAMI EX could develop methods for liquid handling and instrument coordination, tasks such as liquid pipetting and instrument operation each required a separate SAMI method node to be established, displayed, and connected sequentially under the SAMI interface. Taking the plasmid DNA isolation pipeline as an example (Figure 3), SAMI required three nodes to execute steps one and two involving RES buffer transfer, shaker vortexing, and liquid mixing (Figure 3A). In contrast, Biomek 5 could consolidate these three operations into a single node and easily fine-tuned the detailed pipetting conditions such as speed, tip positions, pipetting time, and others. This capability streamlined the pipeline by reducing the number of nodes and enabling clearer visualization of the workflow during method development (Figure 3E). Specifically, the plasmid DNA isolation pipeline contained three major nodes organized according to buffer components defined by the user: one Biomek node for RES buffer processing (asterisk in Figure 3B) and two additional nodes for LYS and NEU buffer treatments (black dots in Figure 3B).

This workflow minimizes human intervention and enhances cost efficiency by automating four major steps: bacterial inoculation, plasmid DNA isolation, purification, and quantification. In addition, the main buffers can be prepared in-house to further improve the cost-effectiveness of the miniprep process. As the automation scale increased to high-throughput levels, consistency became critical for downstream assays such as DNA transfection and PCR. Low DNA yield or high variability in concentration created challenges for concentration normalization. Prolonged incubation of bacteria at 30 °C for 48 hours allowed low-density cultures to catch up without overgrowth of high-density cultures, making the yield less dependent on the initial bacterial density and thus minimizing variation. Based on DNA agarose gel electrophoresis, no visible RNA contamination and only minimal genomic DNA carryover were observed. Given the total vector size of 9.1 kb, the observed migration of the supercoiled DNA at approximately 6 kb was expected. A minor fraction of samples displayed an additional sub-6 kb band, possibly attributable to recombined plasmid species. In general, NanoDrop spectrophotometric analysis revealed that lower DNA yield correlated with decreased 260/230 ratios, likely resulting from ethanol carryover, whereas 260/280 ratios remained stable with minimal protein contamination. Although the NanoDrop is widely used for measuring DNA concentrations in plasmid minipreps, it is not well suited for automated, high-throughput quantification. In addition, its absorbance-based measurements can be inflated by the presence of other nucleic acids, such as single-stranded DNA and RNA, as well as by potential contaminants, including organic solvents and proteins. In contrast, the PicoGreen assay, adaptable to automated, high-throughput workflows, specifically detects double-stranded DNA, resulting in lower measured DNA concentrations compared with the NanoDrop. Nevertheless, we also provide NanoDrop measurements to enable proper comparison of DNA yields from our automated workflow with those obtained from conventional miniprep methods. The high R² value above 0.8 indicates a robust positive correlation between the two methods, demonstrating that our minipreps contain high-quality plasmid composed primarily of double-stranded DNA.

## Acknowledgement

CH and CY were supported by the Sanford Burnham Prebys (SBP) NCI Cancer Center Support Grant P30 CA030199. Research reported in this publication was supported by the SBP Functional Genomics Core through NIH Shared Instrumentation Grant S10 OD036254. We would like to acknowledge the Beckman Coulter team, including Brandon K. Corbin, Kenzo Maetani, Michael D. Moran, Eugene B. Tupas, and Liz Chladny, for their hardware support. We would also like to acknowledge the Beckman software team, including Joshua M. Yoder, Amy V. Gibson, Marc A. Post, Zarina K. Waqar, and Daniel Lynch, for their software support.

## Author contributions

CH, AB, AJD, MJ, and PDA conceptualized and designed the studies. CY, JAY, YW, AG, and CJK participated in methodology development and contributed to the automation workflow establishment. CY and CH performed the experiments and analyzed data. CH, CY, and AB wrote the manuscript.

## Reference

1. Elnagar MA, Ibrahim MF, Albert M, M Talal M, Abdelfattah MM, El-Dabaa E, Helwa R. Homemade plasmid Miniprep solutions for affordable research in low-fund laboratories. AMB Express. 2022 Nov 1;12(1):137. doi: 10.1186/s13568-022-01483-x. PMID: 36319914; PMCID: PMC9626700.

2. Koontz L. Explanatory chapter: how plasmid preparation kits work. Methods Enzymol. 2013;529:23–8. doi: 10.1016/B978-0-12-418687-3.00002-1. PMID: 24011033.

3. Yin JA, Frick L, Scheidmann MC, Liu T, Trevisan C, Dhingra A, Spinelli A, Wu Y, Yao L, Vena DL, Knapp B, Guo J, De Cecco E, Ging K, Armani A, Oakeley EJ, Nigsch F, Jenzer J, Haegele J, Pikusa M, Täger J, Rodriguez-Nieto S, Bouris V, Ribeiro R, Baroni F, Bedi MS, Berry S, Losa M, Hornemann S, Kampmann M, Pelkmans L, Hoepfner D, Heutink P, Aguzzi A. Arrayed CRISPR libraries for the genome-wide activation, deletion and silencing of human protein-coding genes. Nat Biomed Eng. 2025 Jan;9(1):127–148. doi: 10.1038/s41551-024-01278-4. Epub 2024 Dec 4. PMID: 39633028; PMCID: PMC11754104.

4. Farzad MS, Pedersen BM, Mogensen HS, Borsting C. Development of an automated AmpliSeq library building workflow for biological stain samples on the Biomek((R)) 3000. Biotechniques. 2020;68(6):342–4. Epub 2020/03/07. doi: 10.2144/btn-2019-0156. PubMed PMID: 32141765.

5. Van der Heijden S, de Oliveira SJ, Kampmann ML, Borsting C, Morling N. Comparison of manual and automated AmpliSeq workflows in the typing of a Somali population with the Precision ID Identity Panel. Forensic Sci Int Genet. 2017;31:118–25. Epub 2017/09/25. doi: 10.1016/j.fsigen.2017.09.009. PubMed PMID: 28938152.

6. Dawes JC, Webster P, Iadarola B, Garcia-Diaz C, Dore M, Bolt BJ, Dewchand H, Dharmalingam G, McLatchie AP, Kaczor J, Caceres JJ, Paccanaro A, Game L, Parrinello S, Uren AG. LUMI-PCR: an Illumina platform ligation-mediated PCR protocol for integration site cloning, provides molecular quantitation of integration sites. Mob DNA. 2020;11:7. Epub 2020/02/12. doi: 10.1186/s13100-020-0201-4. PubMed PMID: 32042315; PMCID: PMC7001329.

7. Sinha S, Jikare A, Ankulkar R, Mirza Y. Development of miniaturized agar based assays in 96-well microplates applicable to high-throughput screening of industrially valuable microorganisms. J Microbiol Methods. 2022;199:106526. Epub 2022/06/24. doi: 10.1016/j.mimet.2022.106526. PubMed PMID: 35738492.

8. Chen Y, Kaplan Lease N, Gin JW, Ogorzalek TL, Adams PD, Hillson NJ, Petzold CJ. Modular automated bottom-up proteomic sample preparation for high-throughput applications. PLoS One. 2022;17(2):e0264467. Epub 2022/02/26. doi: 10.1371/journal.pone.0264467. PubMed PMID: 35213656; PMCID: PMC8880914.

9. Mardis E, McCombie WR. Preparing Plasmid Subclones for Capillary Sequencing. Cold Spring Harb Protoc. 2017;2017(7):pdb prot094581. Epub 2016/11/03. doi: 10.1101/pdb.prot094581. PubMed PMID: 27803278.

10. Stella S, Vitale SR, Massimino M, Puma A, Tomarchio C, Pennisi MS, Tirro E, Romano C, Martorana F, Stagno F, Di Raimondo F, Manzella L. A Novel System for Semiautomatic Sample Processing in Chronic Myeloid Leukaemia: Increasing Throughput without Impacting on Molecular Monitoring at Time of SARSCoV-2 Pandemic. Diagnostics (Basel). 2021;11(8). Epub 2021/08/28. doi: 10.3390/diagnostics11081502. PubMed PMID: 34441436; PMCID: PMC8391152.

11. Yu C, Caothien R, Jackson M, Nakao B, Pham A, Tam L, Roose-Girma M. Advanced Technologies and Automation in mES Cell Workflow. Methods Mol Biol. 2023;2631:183–206. Epub 2023/03/31. doi: 10.1007/978-1-0716-2990-1_7. PubMed PMID: 36995668.

12. Mardis E, McCombie WR. Preparing Polymerase Chain Reaction (PCR) Products for Capillary Sequencing. Cold Spring Harb Protoc. 2017;2017(7):pdb prot094599. Epub 2016/11/03. doi: 10.1101/pdb.prot094599. PubMed PMID: 27803282.

13. Ohta A, Kawai S, Pretemer Y, Nishio M, Nagata S, Fuse H, Yamagishi Y, Toguchida J. Automated cell culture system for the production of cell aggregates with growth plate-like structure from induced pluripotent stem cells. SLAS Technol. 2023 Dec;28(6):433–441. doi: 10.1016/j.slast.2023.08.002. Epub 2023 Aug 9. PMID: 37562511.

14. Santacruz D, Enane FO, Fundel-Clemens K, Giner M, Wolf G, Onstein S, Klimek C, Smith Z, Wijayawardena B, Viollet C. Automation of high-throughput mRNA-seq library preparation: a robust, hands-free and time efficient methodology. SLAS Discov. 2022;27(2):140–7. Epub 2022/01/31. doi: 10.1016/j.slasd.2022.01.002. PubMed PMID: 35093290.

15. Kind D, Baskaran P, Ramirez F, Giner M, Hayes M, Santacruz D, Koss CK, El Kasmi KC, Wijayawardena B, Viollet C. Automation enables high-throughput and reproducible single-cell transcriptomics library preparation. SLAS Technol. 2022;27(2):135–42. Epub 2022/01/22. doi: 10.1016/j.slast.2021.10.018. PubMed PMID: 35058211.

16. Arrigoni L, Ferrari F, Weller J, Bella C, Bonisch U, Manke T. AutoRELACS: automated generation and analysis of ultra-parallel ChIP-seq. Sci Rep. 2020;10(1):12400. Epub 2020/07/28. doi: 10.1038/s41598-020-69443-8. PubMed PMID: 32709929; PMCID: PMC7381599.

17. Pajak L, Zhang R, Pittman C, Roby K, Boyer S. Automated Genomic and Proteomic Applications on the Biomek® NX Laboratory Automation Workstation. SLAS Technology. 2004;9(3):177–84. doi: 10.1016/j.jala.2004.04.006.

18. Wu Q, Sui X, Tian R. [Advances in high-throughput proteomic analysis]. Se Pu. 2021;39(2):112–7. Epub 2021/07/07. doi: 10.3724/SP.J.1123.2020.08023. PubMed PMID: 34227342; PMCID: PMC9274848.

19. Ball M, Romanovsky E, Schnecko F, Kirchner M, Neumann O, Brandt R, Beck S, Seker-Cin H, Kluck K, Ourailidis I, Goldschmid H, Fink A, Volckmar AL, Menzel M, Allgäuer M, Schirmacher P, Budczies J, Stenzinger A, Kazdal D. Clinical Implementation of a High-Throughput Automated Comprehensive Genomic Profiling Test: TruSight Oncology 500 HT. J Mol Diagn. 2025 Feb;27(2):154–162. doi: 10.1016/j.jmoldx.2024.11.005. Epub 2024 Dec 12. PMID: 39674366; PMCID: PMC12179514.

20. Yang CC, Deshpande AJ, Jackson M, Adams PD, Lynch D, Gibson AV, Waqar ZK, Beketova A, Yin JA, Huang CT. Automation of high-throughput arrayed mammalian cell line cultivation. bioRxiv [Preprint]. 2025 Oct 4:2025.10.03.676043. doi: 10.1101/2025.10.03.676043. PMID: 41256726; PMCID: PMC12621701.

21. Yang CC, Deshpande AJ, Jackson M, Adams PD, Altman Y, Yin JA, Wu Y, Post MA, Beketova A, Huang CT. Automation of high-throughput arrayed lentivirus production and titration. bioRxiv [Preprint]. 2025 Oct 5:2025.10.04.680488. doi: 10.1101/2025.10.04.680488. PMID: 41256661; PMCID: PMC12621811.

22. Yang CC, Deshpande AJ, Jackson M, Adams PD, Pasquale EB, Murad R, Yin JA, Wu Y, Beketova A, Huang CT. Automation of high-throughput workflow for arrayed CRISPR activation library screening. bioRxiv [Preprint]. 2025 Nov 12:2025.11.10.687722. doi: 10.1101/2025.11.10.687722. PMID: 41292928; PMCID: PMC12642482.

